# Characterization of a replicating mammalian orthoreovirus with tetracysteine tagged μNS for live cell visualization of viral factories

**DOI:** 10.1101/174235

**Authors:** Luke D. Bussiere, Promisree Choudhury, Bryan Bellaire, Cathy L. Miller

**Affiliations:** From the Department of Veterinary Microbiology and Preventive Medicine, College of Veterinary Medicine, Iowa State University, Ames, Iowa 50011, USA; Interdepartmental Microbiology Program, Iowa State University, Ames, Iowa 50011, USA

**Author notes:** Corresponding author. Mailing address: Department of Veterinary Microbiology and Preventive Medicine, College of Veterinary Medicine, Iowa State University, 1907 ISU C Drive, VMRI Building 3, Ames, IA 50011.

**Keywords:** Mammalian orthoreovirus, virus factories, tetracysteine tag, live cell imaging, nonstructural μNS protein

## Abstract

Within infected host cells, mammalian orthoreovirus (MRV) forms viral factories (VFs) which are sites of viral transcription, translation, assembly, and replication. MRV non-structural protein, μNS, comprises the structural matrix of VFs and is involved in recruiting other viral proteins to VF structures. Previous attempts have been made to visualize VF dynamics in live cells but due to current limitations in recovery of replicating reoviruses carrying large fluorescent protein tags, researchers have been unable to directly assess VF dynamics from virus-produced μNS. We set out to develop a method to overcome this obstacle by utilizing the 6 amino-acid (CCPGCC) tetracysteine (TC)-tag and FlAsH-EDT2 reagent. The TC-tag was introduced into eight sites throughout μNS, and the capacity of the TC-μNS fusion proteins to form virus factory-like (VFL) structures and colocalize with virus proteins was characterized. Insertion of the TC-tag interfered with recombinant virus rescue in six of the eight mutants, likely as a result of loss of VF formation or important virus protein interactions. However, two recombinant (r)TC-μNS viruses were rescued and VF formation, colocalization with associating virus proteins, and characterization of virus replication were subsequently examined. Furthermore the rTC-μNS viruses were utilized to infect cells and examine VF dynamics using live cell microscopy. These experiments demonstrate active VF movement with fusion events as well as transient interactions between individual VFs, and demonstrate the importance of microtubule stability for VF fusion during MRV infection. This work provides important groundwork for future in depth studies of VF dynamics and host cell interactions.

**Importance:** MRV has historically been used as a model to study the double-stranded RNA (dsRNA) *Reoviridae* family, which infect and cause disease in humans, animals, and plants. During infection, MRV forms VFs that play a critical role in virus infection, but remain to be fully characterized. To study VFs, researchers have focused on visualizing the non-structural protein μNS which forms the VF matrix. This work provides the first evidence of recovery of replicating reoviruses in which VFs can be labeled in live cells via introduction of a TC-tag into the μNS open reading frame. Characterization of each recombinant reovirus sheds light on μNS interactions with viral proteins. Moreover, utilizing the TC labeling FlAsH-EDT2 biarsenical reagent to visualize VFs, evidence is provided of dynamic VF movement and interactions at least partially dependent on intact microtubules.

## Introduction

Mammalian orthoreovirus (MRV) is a segmented, double-stranded RNA (dsRNA) virus that has been utilized as a tool to study the life cycle of the virus family *Reoviridae*. The virus is also of particular interest as it is classified as an oncolytic virus and is currently being studied in various pre-clinical and clinical trials to treat multiple tumor types (1). During MRV infection, inclusion bodies called viral factories (VFs) form, which were first identified in 1962 and later described as small particles that coalesced to form a reticulum-like structure around the nucleus (2, 3). VFs have historically been hypothesized to be the location of viral replication and assembly of viral core particles (4, 5), and recent data showing core particle recruitment and positive strand RNA synthesis at VFs (6, 7) and translation of viral RNA within and around VFs (8) suggest they are also the site of viral transcription and translation.

VFs are largely comprised of the viral non-structural protein μNS which is suggested to be the only virus protein required to form the matrix of VFs (9). When expressed in transfected cells from a plasmid, μNS forms inclusions similar to VFs that are termed viral factory-like (VFL) structures (9). Moreover, μNS formation of VFLs results in specific VFL localization of each of the five viral structural proteins that make up the core particle (λ1, λ2, λ3, σ2, and μ2), the non-structural σNS protein that is involved in virus translation and replication, as well as the intact core particle itself (6, 9-11). The mechanism of VFL formation by μNS is not fully understood, however, several regions of the protein, including the carboxyl (C)-terminal 7 amino-acids (AA), and a putative metal chelating structure formed by AA His570 and Cys572 have been shown to be necessary for VFL formation in transfected cells and replication in infected cells (12-14). Additionally, the C-terminal 250 amino-acids (AA 471-721) of μNS, which comprise a predicted coiled-coil domain, are sufficient to form VFL structures (12, 15). Deletion of AA 471-561 disrupts VFL formation, suggesting this region also plays a role in the assembly of these structures (12).

Much of the work defining VFs has been done using transient transfection of plasmids expressing μNS and other virus proteins, primarily because it has thus far been difficult to visualize these proteins during infection in live cells. While researchers have been able to add fluorescent tags to viral proteins in other viruses and recover recombinant virus, the MRV genome has only recently been made amenable to reverse genetics technologies (16-19). Utilizing this technology, a virus in which the coding region within the S4 gene segment was replaced by the EGFP gene has been recovered, however, this virus replicates only when propagated in cells expressing the S4 gene product σ3 (16). Both replicating and non-replicating recombinant viruses have also been recovered with fluorescent proteins iLOV and UnaG independently expressed downstream of the N-terminal half of viral spike protein, σ1 (σ1-N), as well as a fusion of σ1-N with UnaG which could infect cells but was unable to replicate in the absence of wildtype MRV (19, 20). In addition, recombinant reoviruses in which small protein coding sequences (6His, HA-tag, and 3HA-tag) were added to the σ1 protein have been recovered (21). However, there is currently no other published evidence of a replicating, recombinant virus in which μNS or other viral proteins have been successfully tagged with fluorescent or other tags. To overcome this difficulty visualizing VFs in infected live cultures, cells transiently expressing EGFP-μNS have been infected with intermediate subvirion particles (ISVPs) (8), and VFs formed by the dual expression of virus expressed μNS and EGFP-μNS have been examined. This has led to important findings including demonstrating the interaction of the cellular vesicular network with VFs during infection; however, the addition of the non-wildtype EGFP-μNS may alter VF function and virus replication complicating interpretation of results. The ability to rescue a replicating recombinant virus expressing a fluorescent or fluorescent competent tagged μNS would allow for direct visualization of VF dynamics in MRV infected cells.

One possible solution for tagging the μNS protein within the viral genome is to exploit the 4,5–bis(1,3,2-dithioarsolan-2-yl)fluorescein, also known as the fluorescein arsenical helix binder, bis-EDT adduct (FlAsH-EDT2) which fluoresces green when bound to a small 6 AA tag (CCXXCC) (22). FlAsH-EDT2 was first described in 1998 and was shown to fluoresce when bound to two pairs of cysteines separated by two amino-acids (CCXXCC) referred to as a tetracysteine (TC)-tag (22). This technique has been utilized in several recombinant viruses including HIV-1 to visualize TC-Gag protein during infection, and in bluetongue virus, a member of the *Reoviridae* family, to label viral protein VP2 to visualize virus particle entry (23, 24). Once the TC-tag is added to a protein, FlAsH-EDT2, which contains two 1,2-ethanedithiol motifs, creates covalent bonds with the TC motif resulting in fluorescence. FlAsH-EDT2 can be added at any time post infection (p.i.) allowing for visualization of TC-tagged proteins throughout the virus life cycle. As CCPGCC is rare to find within natural proteins and is the preferred FlAsH-EDT2 binding sequence (25), it is a good AA sequence to use for the TC-tag, resulting in highly specific FlAsH-EDT2 binding with little background fluorescence.

In this study we introduced the CCPGCC TC-tag in frame into eight sites throughout the μNS protein and examined the impact of the insertion on known μNS functions. We further demonstrated that we could rescue recombinant viruses expressing the TC-tag at two sites within μNS. Recombinant (r)TC-μNS viruses were characterized with regard to replication, VF formation, and stability of the TC-tag within the M3 genome segment over multiple viral passages. Finally, rTC-μNS viruses were used to examine μNS and VF dynamics in live cells, and to observe the role of microtubules in VF fusion and movement throughout the cell.

## Materials and methods

### Cells, viruses, antibodies, and reagents

CV-1, Vero, and BHK-T7 cells (26) were maintained in Dulbecco’s modified Eagles’s medium (DMEM) (Invitrogen Life Technologies) supplemented with 10% or 2% (Vero during virus amplification) fetal bovine serum (Atlanta Biologicals), penicillin (100 I.U./ml) streptomycin (100 μg/ml) solution (Mediatech), and 1% MEM non-essential amino-acids solution (HyClone).To maintain the T7 polymerase 1 mg/ml of G418 (Alexis Biochemical) was added every fourth passage to BHK-T7s. L929 cells were maintained in Joklik modified minimum essential medium (Sigma-Aldrich) supplemented with: 2% fetal bovine serum, 2% bovine calf serum (HyClone), 2 mM L-glutamine (Mediatech), and penicillin (100 I.U./ml) streptomycin (100 μg/ml) solution. Our laboratory stock of MRV strain type 1 Lang (T1L) originated from the laboratory of B. N. Fields. The virus was propagated and purified as previously described (27). Primary antibodies used were as follows: monoclonal mouse anti-FLAG (α-FLAG) antibody (Sigma Aldrich, F1804), monoclonal mouse α-HA antibody (ThermoFisher, 26183), polyclonal mouse α-μNS, polyclonal rabbit α-μNS, α-μ2, and T1L α-virion antibodies (9, 27-29), monoclonal mouse α-σNS (3E10) and α-λ2 (7F4) antibodies deposited in the Developmental Studies Hybridoma Bank (DSHB) by Dermody, T.S. (30, 31), and rabbit α-MRV(core) (32). Secondary antibodies used are Alexa 594- and 488-conjugated donkey anti-mouse or anti-rabbit IgG antibodies (Invitrogen Life Technologies). FlAsH-EDT2 (ThermoFisher, Cayman Chemical) was used at a final concentration of 2.5 μM. Nocodazole (Acros Organics) was used at a final concentration of 10 μM.

### Plasmid construction

pCI-σNS, pCI-λ1, pCI-λ2, pCI-λ3/HA, and pCI-σ2 were previously described (6, 10, 11). The T1L MRV reverse genetics plasmids pT7-M1-S1-S2-S4-RZ, pT7-L3-S3-RZ, pT7-L1-M2-RZ, pT7-L2 and pT7-M3 were previously described (16, 17). The Flag-tagged μ2 plasmid (pCI-Flag/M1 T3D^C^) was made by PCR amplification of a plasmid encoding the Flag-tag and the M1 5’ end (nts 1-564) using forward and reverse primers (IDT) containing an XhoI site and an EcoRV site respectively, flanking Flag-M1 gene sequence homology. The PCR product and pCI-μ2 (33) were digested with XhoI and EcoRV and ligated. The TC-tagged μNS plasmids were constructed utilizing synthetic dsDNA gBlocks (IDT) containing each of the described mutations and the surrounding μNS sequence flanked on each end by restriction sites as follows: TC-μNS(#1) (SacI-BbvCI), TC-μNS(#2) (BbvCI-XmaI), TC-μNS(#4) and TC-μNS(#5) (BclI-BstEII), TC-μNS(#6) and TC-μNS(#7) (BstEII-SalI), TC-μNS(N-term) (SacI-PciI), and TC-μNS(C-term) (SalI-NotI). Each gBlock and pT7-M3 were digested with the indicated restriction enzymes, and vector and insert fragments were ligated. All restriction enzymes were purchased from NEB.

### Transfection, infection and reverse genetics

For transfections and infections, BHK-T7 and CV-1 cells were seeded at a concentration of 2.0×10^5^ and 1.0×10^5^ cells per well, respectively, in 12-well plates containing 18 mm glass coverslips the day before transfection or infection. For transfection, 1 μg of plasmid DNA and 3 μl of TransIT-LT1 reagent (Mirus Bio) were added to 100 μl of Opti-MEM reduced serum medium (Invitrogen Life Technologies), incubated at room temperature for 30 mins, added dropwise to cells, and incubated at 37°C overnight. CV-1 cells were infected with T1L, rTC-μNS(C-term-T1L)/passage 5 (P5), or rTC-μNS(#7-T1L)/P5 at an MOI = 1 for 1 h with shaking and then incubated at 37°C overnight. For reverse genetics, 5.0×10^5^ BHK-T7 cells were plated on a 6-well plate (9.5 cm^2^) (Corning Inc.) and transfected as previously described (27). The cells were incubated for six days at which point cells and media were subjected to three freeze/thaw cycles, and standard L929 cell plaque assays were performed (34). Recovered plaques were passaged ten times on Vero cells for 7-28 days to allow for replication of slow growing virus.

### Immunofluorescence assay

18 h post-transfection (p.t.) or p.i., cells were treated or not with FlAsH-EDT2 diluted to 2.5 μM in Opti-MEM (Invitrogen Life Technologies) for 45-90 mins then fixed with 4% paraformaldehyde for 20 mins, and washed twice with phosphate buffered saline (PBS) (137 mM NaCl, 3 mM KCl, 8 mM Na_2_HP0_4_, 1.5 mM KH_2_PO_4_, pH 7.4). Cells were permeabilized with 0.2% Triton X-100 in PBS for 5 mins, washed twice with PBS, and blocked with 1% bovine serum albumin in PBS (PBSA) for 10 mins. Cells were then incubated for 45 min at room temperature with primary antibodies diluted in PBSA and washed two times with PBS, followed by incubation with secondary antibody diluted in PBSA for 30 mins and two additional PBS washes. Labeled cells were mounted with ProLong Gold antifade reagent with DAPI (4, 6-diamidino-2-phenylindole dihydrochloride) (Invitrogen Life Technologies) on slides. Each coverslip was then examined on a Zeiss Axiovert 200 inverted microscope equipped with fluorescence optics. Representative pictures were taken by a Zeiss AxioCam MR colour camera using AxioVision software (4.8.2). Plot profiles were utilized using ImageJ (2.0.0-rc-49/1.51d) to determine pixel intensity of viral protein localization relative to μNS-labeled VFLs to compare wildtype μNS to TC-μNS. A significant difference (*p* < 0.05), denoted as a (-) in Table 1, was determined using JMP Pro (12.0.1). Images were prepared using Adobe Photoshop and Illustrator software (Adobe Systems).

**Table 1.** Summary of TC-μNS mutant properties. Each TC-μNS mutant, location of each CCPGCC tag, VFL formation, FlAsH-EDT2 labeling, virus recovery, and MRV protein colocalization compared to wildtype μNS used in this study is summarized within this table. A negative symbol (-) denotes there was no VFL formation and no FlAsH-EDT2 labeling, or that there was a significantly diminished (*p* < 0.05) colocalization of TC-μNS and interacting proteins compared to wildtype μNS. A positive symbol (+) denotes there was VFL formation and FlAsH-EDT2 labeling, or that there was no significant decrease in colocalization of μNS and interacting proteins compared to wildtype μNS. ND=not determined.

| TC-μNS | Location of CCPGCC | VFL formation | FlAsH-EDT2 labeled | Virus recovery | Protein colocalization compared to wildtype μNS |  |  |  |  |  |
| --- | --- | --- | --- | --- | --- | --- | --- | --- | --- | --- |
|  |  |  |  |  | σNS | μ2 | λ1 | λ2 | σ2 | λ3 |
| #1 | AA 28 | + | + | No | - | + | + | - | + | + |
| #2 | AA 132 | - | - | No | ND | ND | ND | ND | ND | ND |
| #4 | AA 334 | + | + | No | + | + | - | - | + | - |
| #5 | AA 491 | - | - | No | ND | ND | ND | ND | ND | ND |
| #6 | AA 614 | + | + | No | + | + | - | + | + | - |
| #7 | AA 545 | + | + | Yes | + | - | - | + | + | - |
| N-term | N-terminus | + | + | No | - | + | + | + | + | + |
| C-term | C-terminus | + | + | Yes | + | + | + | + | + | + |

### Replication assay

L929 cells were seeded at a concentration of 2.5×10^6^ cells per 60 mm dish (Corning Inc.). 24 h post-seeding, cells were infected with wildtype T1L, rTC-μNS(C-term-T1L)/P2 or /P7, or rTC-μNS(#7-T1L)/P2 virus at an MOI of 0.2. At 0, 12, 24, 36, and 48 h p.i. cells were harvested and subjected to three freeze/thaw cycles, then standard MRV plaque assays were performed on L929 cells to determine viral titers (34).

The mean and standard deviation were determined from two experimental replicates in two different biological replicates. A student’s t-test was used to determine the *p*-value using Microsoft Excel (Microsoft Office). After the viral titer was determined 8% paraformaldehyde was added to each well and left to incubate overnight. The next day the paraformaldehyde and overlay was removed, and cells were washed with PBS twice, and 1% crystal violet in 20% ethanol was added and incubated at room temperature for 15 mins. Cells were washed twice with water, and the plaques were imaged using a ChemiDoc XRS Imaging System (Bio-Rad). Plaque size was then determined by measuring five plaques from each virus using ImageJ (2.0.0-rc-49/1.51d) and the average was calculated. Images were prepared using Adobe Photoshop and Illustrator (Adobe Systems).

### Immunoblot

L929 cells were plated at a concentration of 1×10^6^ cells per well in a 6-well plate. 24 h post-seeding, cells were mock infected or infected with T1L, rTC-μNS(C-term-T1L)/P5, or rTC-μNS(#7-T1L)/P5 at an MOI = 0.25. At 0, 12, 24 and 48 h p.i. cells were washed twice with PBS and collected in 2X protein loading buffer (125 mM Tris-HCl pH 6.8, 200 mM dithiothreitol (DTT), 4% sodium dodecyl sulfate (SDS), 0.2% bromophenol blue, 20% glycerol). Alternatively, 5×10^5^ PFUs of each virus was diluted to the same volume using PBS + 2 mM MgCl_2_ and then 2X protein loading buffer was added to each sample. Cell lysates/virions were separated by SDS-polyacrylamide gel electrophoresis (SDS-PAGE) and then transferred to nitrocellulose by electroblotting. Nitrocellulose membranes were incubated with primary and secondary antibodies in Tris buffered saline (20 mM Tris, 137 mM NaCl, pH7.6) with 0.25% Tween 20 (TBST) for 18 h and 4 h respectively, followed by addition of PhosphaGLO AP (SeraCare) substrate and imaging on a ChemiDoc XRS Imaging System (Bio-Rad).

### MRV genome analysis and reverse transcription

1.0×10^8^ PFUs of T1L, rTC-μNS(C-term-T1L)/P2, /P5, /P7, or /P10, or rTC-μNS(#7-T1L)/P2 or /P5 and 1.0×10^9^ PFUs of T1L (T1L 10X) were subjected to TRIzol LS (Life Technologies) extraction via manufacturer’s instructions. Briefly viruses were homogenized in TRIzol LS and the addition of chloroform separated each sample into proteins, DNA, and RNA phases. The RNA phase was collected and isopropanol and ethanol were added to precipitate and wash RNA. Extracted RNA was separated on 10% SDS-PAGE at a constant 20 mA for 12 hrs, and the gel was incubated in water with 3X gel red (Phenix Research Products) for 1 h and imaged on a ChemiDoc XRS Imaging System (Bio-Rad). Extracted RNA was also subjected to reverse transcription (RT)-PCR using SuperScript IV (Invitrogen Life Technologies) as per manufacturer’s instructions for sequencing.

### Live cell imaging of VFs

BHK-T7 and Vero cells were seeded on a 12 well, 14 mm glass bottom plate (MatTek Corporation) at a concentration of 2×10^5^ and 7.5×10^4^ cells per well, respectively, and then infected the following day with T1L at an MOI of 100, rTC-μNS(C-term-T1L)/P2 at an MOI of 5 (BHK-T7 cells), or rTC-μNS(C-term-T1L)/P5 at an MOI of 100 (Vero cells). For BHK-T7 cells, at 18 h p.i., media was removed and cells were washed twice with DMEM without phenol red (HyClone) supplemented with penicillin (100 I.U./ml), streptomycin (100 μg/ml) solution, 1% MEM non-essential amino-acids solution, and 25 mM HEPES (4-(2-Hydroxyethyl)piperazine-1-ethanesulfonic acid), followed by addition of FlAsH-EDT2 diluted to 2.5 μM in DMEM and incubated at 37^o^C with shaking every 15 mins for a total of 45 mins, at which point 800 μl of DMEM was added to each well and imaging was initiated. For Vero cells, at 10 h p.i. cells were washed and treated with FlAsH-EDT2 until 12 h p.i. at which point FlAsH-EDT2 was removed, media was replenished and imaging was initiated. In experiments involving nocodazole, at 6 h p.i. 10 μM nocodazole was added and maintained throughout the experiment. All cells were examined using an Olympus IX071 inverted fluorescence microscope on a vibration table, equipped with an environmental control chamber heated to 37°C. Still images and video were captured through a 40x apochromatic objective by a high resolution Hamamatsu CCD camera using MetaMorph for Olympus MetaMorph Advanced (V 7.7.7.0). All image exposure conditions were maintained throughout the experiment and background levels were set using FlAsH-EDT2 expression in wildtype T1L infected cells. Still images were processed using ImageJ (2.0.0-rc-49/1.51d) and assembled for publication using Adobe Photoshop and Illustrator (Adobe Systems).

## Results

### Construction of plasmids expressing the TC-tag from within the μNS protein

Specific regions within the μNS protein have previously been identified that are required for recruitment of the viral core and six other MRV proteins to VFs, and for forming VFs (6, 7, 9, 10, 12) (Fig. 1A). As our goal was to rescue a type 1 Lang (T1L) MRV virus expressing the TC-tag, CCPGCC, from within the μNS protein, we reasoned that there would be sites of insertion that would not allow rescue of virus as a result of interference with μNS functions during infection. For this reason, to locate a position within the protein that would be tolerated during virus infection, we inserted the nucleotide sequence encoding CCPGCC at eight positions throughout a μNS-encoding M3 gene expression plasmid. We initially reasoned that utilization of existing amino-acids of the TC-tag from within the μNS protein may prevent disruption of μNS function and constructed clones in which we incorporated the remainder of the tag into an existing PG, by flanking the PG residues with two CC residues at AA 28 [TC-μNS(#1)], 132 [TC-μNS(#2)] and 334 [TC-μNS(#4)], or CP, by flanking the CP with a C and GCC at AA 491 [TC-μNS(#5)]. Although μNS is highly conserved between strains, to improve our chance of success in rescuing a recombinant virus expressing a tagged μNS, we performed a sequence comparison between the μNS protein from the three major MRV serotypes (T1L, T2J, and T3D), and identified two areas of 6 AA that were not highly conserved. We replaced these regions with CCPGCC in the T1L μNS (TC-μNS(#6)- AA 614 EAAAKC to CCPGCC, and TC-μNS(#7)-AA 545 QSLNAQ to CCPGCC). Finally, we reasoned we may be able to rescue a virus where the μNS protein sequence was not disrupted and the CCPGCC was added as a fusion to either the μNS amino (N)- or carboxyl (C)-terminus to make TC-μNS(N-term) and TC-μNS(C-term) (Fig. 1A).

**Figure 1.**
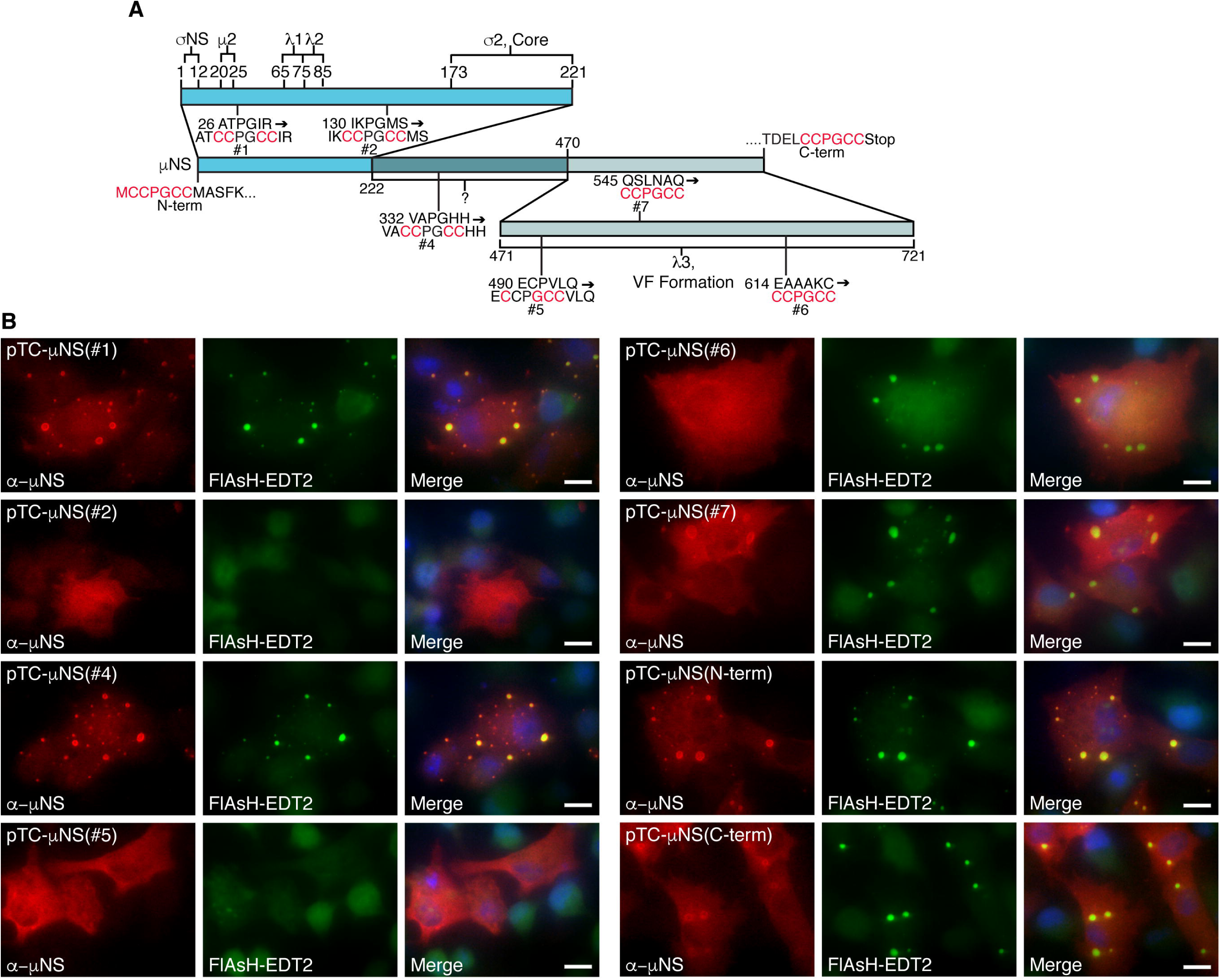
FlAsH-EDT2 labeling and VFL formation in μNS CCPGCC mutants. (A) A diagram of μNS depicting the location and sequence of TC-μNS mutations and previously described functional regions of μNS necessary for protein localization. (B) BHK-T7 cells were transfected with indicated plasmids, and at 18 h p.t., cells were labeled with FlAsH-EDT2 (middle column) for 90 min, then fixed and immunostained with α-μNS antibody (left column), followed by Alexa 594-conjugated donkey antirabbit IgG. A merged image is also shown with DAPI staining (right column). Images are representative of the observed phenotype. Bars = 10 μm.

### VFL formation and biarsenical labeling of TC-μNS in transfected cells

Insertion of the TC-tag does not ensure the protein will be labeled upon exposure to biarsenical compounds as folding of the tagged protein may prevent or interfere with compound binding. It is also plausible that the TC-tag would prevent VFL formation due to a conformational change in μNS resulting in loss of μNS-μNS association. In order to examine FlAsH-EDT2 labeling and VFL formation of the TC-μNS fusion proteins, we transfected BHK-T7 cells with plasmids expressing each of the eight TC-tagged proteins. At 18 h p.t., FlAsH-EDT2 reagent was added. We began to observe fluorescence within 15 mins in most samples (data not shown), that increased in intensity throughout the labeling period of 90 mins, at which point the FlAsH-EDT2 reagent was removed and cells were fixed and subjected to immunofluorescence assays with antibodies against μNS, followed by Alexa 594-conjugated secondary antibodies (Fig. 1B). Upon microscopic observation, we found colocalization of the FlAsH-EDT2 and μNS staining (Fig. 1B) indicating that the FlAsH-EDT2 was binding TC-μNS in six of the eight proteins, including TC-μNS(#1), TC-μNS(#4), TC-μNS(#6), TC-μNS(#7), TC-μNS(N-term), and TC-μNS(C-term). TC-μNS(#5) exhibited weak FlAsH-EDT2 labeling, and TC-μNS(#2) did not exhibit detectable labeling, although there were high levels of μNS protein expressed in these cells. We next examined the effect of the TC-tag on mutant VFL formation and we found that TC-μNS(#1), TC-μNS(#4), TC-μNS(#6), TC-μNS(#7), TC-μNS(N-term), and TC-μNS(C-term) formed VFLs. TC-μNS(#1), TC-μNS(#4), and TC-μNS(N-term) had little to no diffuse μNS staining and formed subjectively tighter VFLs than TC-μNS(#6), TC-μNS(#7), and TC-μNS(C-term) which formed less distinct VFLs with varying levels of diffuse μNS staining. TC-μNS(#2) and TC-μNS(#5) did not form VFLs and instead were entirely diffuse in all transfected cells. These results suggest that the TC-tag can be added at multiple sites throughout the μNS protein without disrupting VFL formation and that the expressed TC-μNS protein was label-competent utilizing FlAsH-EDT2.

### Insertion of the TC-tag within the N-terminal two thirds of μNS leads to diminished recruitment of viral proteins to VFLs

We hypothesized that we could also utilize these mutants to learn more about μNS localization with MRV proteins by investigating the interaction of each mutant with individual virus proteins that were previously identified as μNS associating partners. We separated the TC-μNS mutants based on previously identified functions of μNS to include the N-terminal third (AA 1-221), the central third (AA 222-470), and the C-terminal third (AA 471-721). Previous studies have found that the μNS N-terminal third is both necessary and sufficient to bind virus proteins σNS, μ2, λ1, λ2, and σ2, the C-terminal third is necessary and sufficient to bind λ3, form VFs, and bind cellular clathrin, and the central third, which has been implicated in recruiting cellular protein Hsc70 to VFs (6, 7, 9, 10, 12, 35, 36). To examine the impact of the introduced TC-μNS mutations we co-transfected cells with each pTC-μNS mutant or wildtype pT7-M3 along with plasmids expressing each of the other six virus proteins individually. As TC-μNS(#2) and TC-μNS(#5) did not form VFLs, we did not examine the impact of these mutations on μNS localization with other viral proteins. At 18 h p.t., cells were fixed and stained with antibodies against μNS and each respective protein or associated protein tag followed by Alexa 594- and 488-conjugated secondary antibodies. Three representative pictures of each condition were acquired and μNS recruitment of λ1, λ2, λ3, σ2, and σNS to VFLs, or μ2 recruitment of μNS to microtubules was quantified by comparing the pixel intensity of μNS and associating proteins at VFLs or microtubules. Each quantified interaction of TC-μNS and associating protein was made relative to wildtype μNS and a student’s t-test was used to determine if each TC-μNS mutant was significantly decreased, (*p* < 0.05) denoted by a (-), from wildtype (Table 1). We found that each TC-μNS mutant had a statistically significant diminished colocalization with one or more viral proteins with the exception of the C-terminal mutant which maintained wildtype levels of colocalization of all proteins. Since the N-terminal third of μNS is necessary for multiple interaction with μNS associating proteins, it is not surprising that the insertion within this coding region resulted in disruption of colocalization of the most proteins. As the central and C-terminal thirds of μNS are not implicated in viral protein recruitment to VFLs aside from λ3, it unclear why TC insertions in these regions result in decreased colocalization, but may suggest loss of proper μNS folding in these mutants. Altogether, these findings suggest that the C-terminus may be the most amenable region of μNS for TC-tag insertion.

### Rescue of MRV expressing TC-μNS

All of the TC-μNS mutants were tested in a plasmid based reverse genetics approach to attempt to rescue viruses containing the CCPGCC motif in the μNS protein (17). Briefly, a modified version of the previously described T7-driven, 4 and 10 plasmid systems was used (17). Three plasmids from the four-plasmid system were used to provide 8 of the MRV genes (pT7-M1-S1-S2-S4, pT7-L3-S3, pT7-L1-M2). The L2 and M3 wildtype and M3 TC mutant genes were each provided from individual plasmids (pT7-L2, pT7-M3/TC-M3 mutants). Plasmids were co-transfected into BHK-T7 cells, and incubated for six days at which point cells and media were harvested and subjected to three freeze/thaw cycles, followed by standard MRV plaque assay on L929 cells (34). Three experiments were done for each mutant, and wildtype M3 and no M3 plasmid positive and negative controls were included in each experiment. To allow for slow growing viruses, plaque assays were incubated for five days and monitored each day for plaques. In each experiment there were greater than 10^4^ plaques formed in the positive control and no plaques formed in the negative control. There were also no plaques recorded in any experiment with pTC-μNS(N-term), pTC-μNS(#1), pTC-μNS(#2), pTC-μNS(#4), pTC-μNS(#5) or pTC-μNS(#6). However, we observed plaques forming from days 3-5 on a single experiment with pTC-μNS(#7) and pTC-μNS(C-term), suggesting we had recovered rTC-μNS(#7-T1L) and rTC-μNS(C-term-T1L) viruses with TC-tagged μNS. Plaques were picked and the viruses were passaged ten times on Vero cells to amplify the viruses. After two passages, CV-1 cells were infected with recombinant viruses and were labeled with FlAsH-EDT2 18 h p.i. and subsequently prepared for immunofluorescence to label μNS. We observed FlAsH-EDT2 labeling of μNS and VFs in cells infected with each virus (Fig. 2) indicating we had recovered two fluorescent competent, recombinant viruses. While both viruses exhibited FlAsH-EDT2 labeling, rTC-μNS(#7-T1L) factories were qualitatively less fluorescent than rTC-μNS(C-term-T1L) factories.

**Figure 2.**
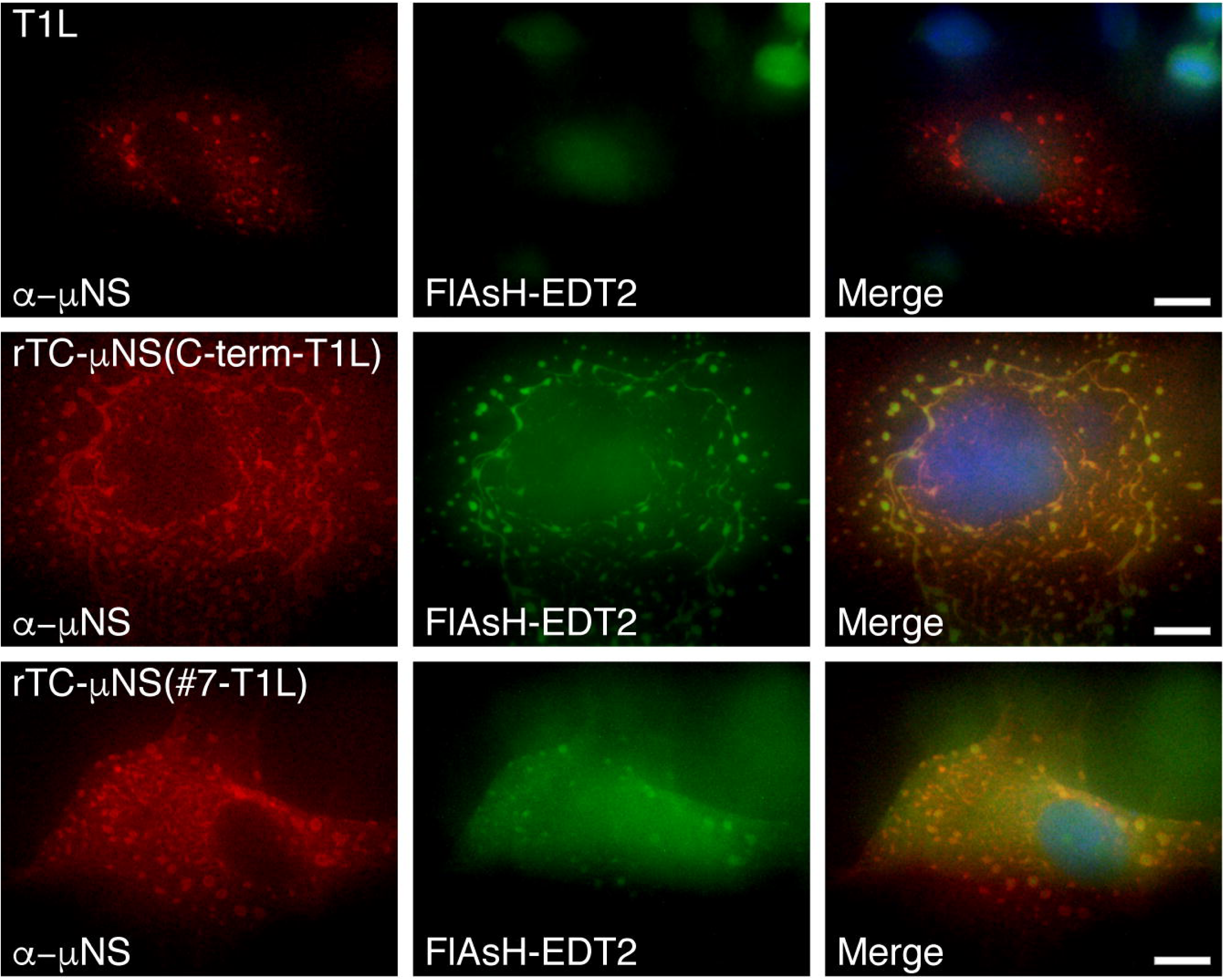
VFs from recombinant viruses label with FlAsH-EDT2. CV-1 cells were infected with T1L, rTC-μNS(C-term-T1L)/P2, or rTC-μNS(#7-T1L)/P2, and at 18 h p.i., cells were labeled with FlAsH-EDT2 (middle column) for 45 min, then fixed and immunostained with α-μNS antibody (left column), followed by Alexa 594-conjugated donkey anti-rabbit IgG. A merged image is also shown with DAPI staining (right column). Images are representative of the observed phenotype. Bars = 10 μm.

### TC-μNS containing viruses exhibit growth deficiencies

After two passages of both rTC-μNS(#7-T1L) and rTC-μNS(C-term-T1L) viruses were titered on L929 cells and replication assays were performed to determine the fitness of the recombinant viruses relative to wildtype T1L. Following infection with an MOI of 0.2, samples were taken at 0, 12, 24, 36, and 48 h p.i., freeze/thawed three times, and subjected to plaque assays on L929 cells (Fig. 3A). While both recombinant viruses replicated comparable to wildtype up to 12 h p.i. we observed a statistically significant attenuation of virus replication at 24, 36, and 48 h p.i. in the recombinant viruses relative to wildtype. Overall the wildtype virus replicated to between 10- and 100-fold higher than rTC-μNS(#7-T1L) and rTC-μNS(C-term-T1L) after 24 h p.i. Despite this attenuation, recombinant viruses were able to replicate to about 10^3^ fold relative to time zero at 48 h p.i., suggesting they were capable of completing the entire virus life cycle, albeit slower than wildtype T1L (Fig. 3A). We additionally measured the plaque size of each virus and found that rTC-μNS(#7-T1L) and rTC-μNS(C-term-T1L) plaques were 55% and 45% the size of wildtype plaques, respectively (Fig. 3B). These findings suggest that addition of the TC-tag in the C-terminal third of μNS inhibits viral infection relative to wildtype virus, however, both rTC-μNS(#7-T1L) and rTC-μNS(C-term-T1L) replicated to titers that should be sufficient for live cell imaging.

**Figure 3.**
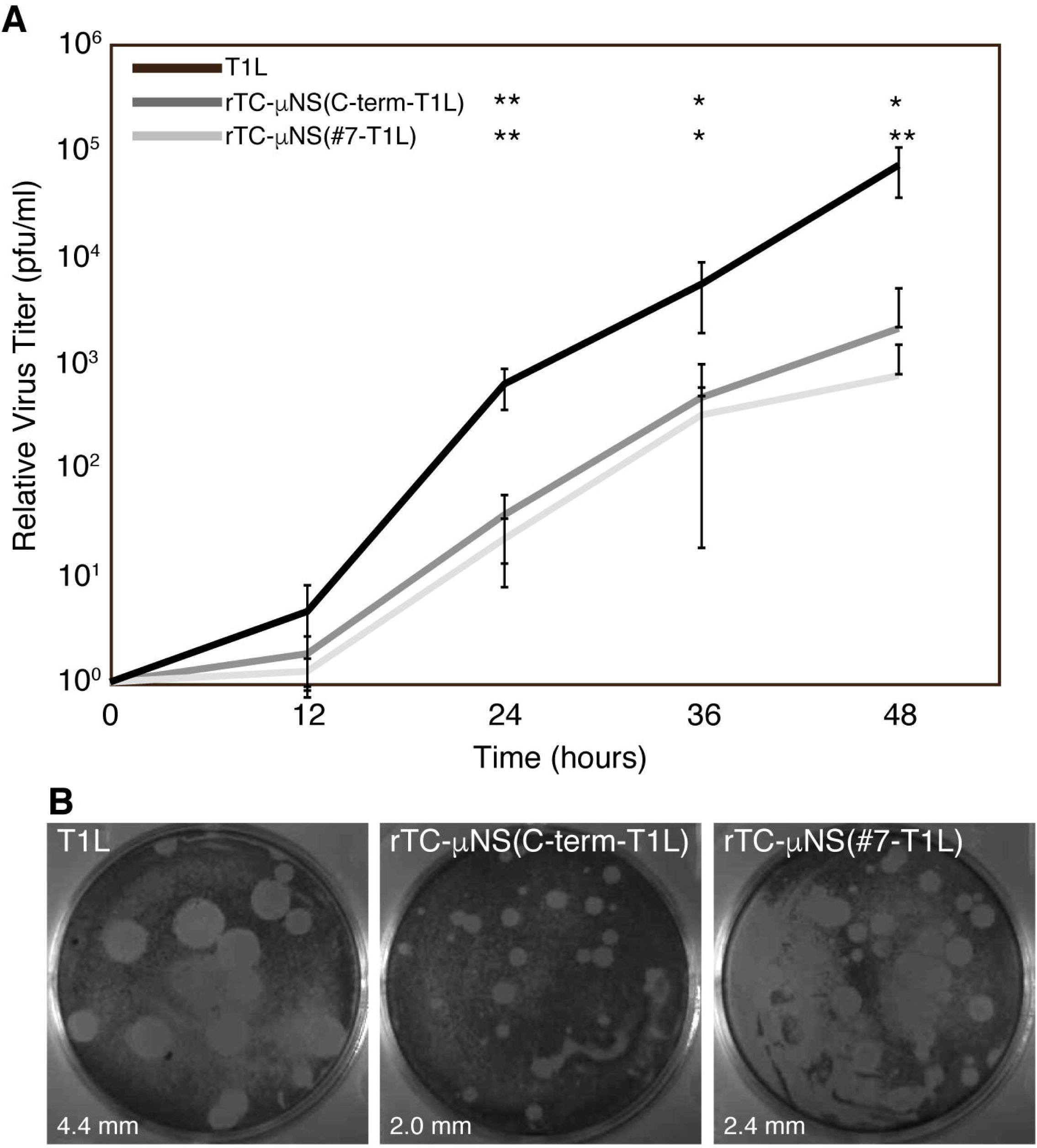
Recombinant TC-μNS virus replication. L929 cells were infected with T1L, rTC-μNS(C-term-T1L)/P2, or rTC-μNS(#7-T1L)/P2 and at 0, 12, 24, 36, and 48 h p.i. cells were harvested. Harvested cells were subjected to standard MRV plaque assay. (A) Plaques from each time point were counted and the relative viral titer increase from time zero is plotted. The means and standard deviations were calculated from two experimental replicates within two different biological replicates. A two-tailed student’s t-test was used to calculate a *p-*value for the significant difference between recombinant virus and wildtype virus in Microsoft Excel, *p* < 0.05 is denoted by an asterisks (*) and *p* < 0.01 is denoted by two asterisks (**). (B) Cells were fixed and stained with crystal violet and imaged to visualize plaque size relative to wildtype T1L. Five plaques were measured using ImageJ in each panel to determine average plaque diameter (bottom left).

### Recombinant viruses retain recruitment of μNS associating proteins to VFs

In an attempt to explain the attenuated nature of rTC-μNS(#7-T1L) and rTC-μNS(C-term-T1L) we examined μNS recruitment of viral proteins to VFs in cells infected with each recombinant virus. CV-1 cells were infected with T1L, rTC-μNS(C-term-T1L)/P5, or rTC-μNS(#7-T1L)/P5 at an MOI of 1 and at 18 h p.i. cells were fixed and stained with antibodies against μNS and σNS, λ2, μ2, or the core particle followed by Alexa 594- and 488-conjugated secondary antibodies (Fig. 4A-C). The core antibody has been shown to bind λ1, λ2, and σ2 (6), and we do not have access to an antibody that specifically detects λ3. We observed recruitment of σNS, λ2, and the core proteins to VFs and recruitment of VFs to microtubules in both recombinant viruses, with rTC-μNS(C-term-T1L) displaying a qualitatively more similar phenotype to wildtype virus compared to rTC-μNS(#7-T1L) (Fig. 4), which consistently possessed somewhat smaller VFs than T1L. This data suggests that rTC-μNS(C-term-T1L) may better depict natural VFs in live cell imaging.

**Figure 4.**
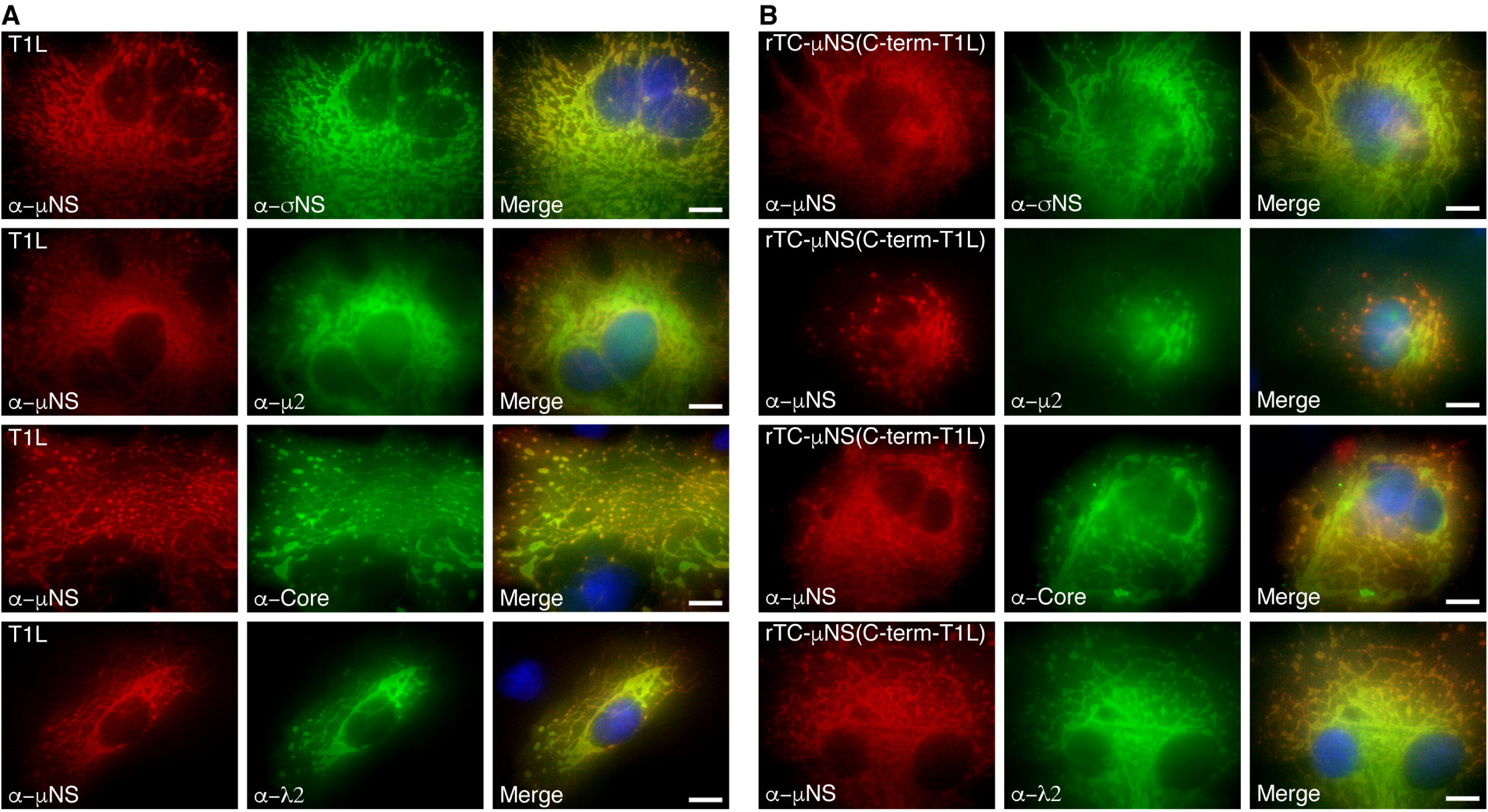
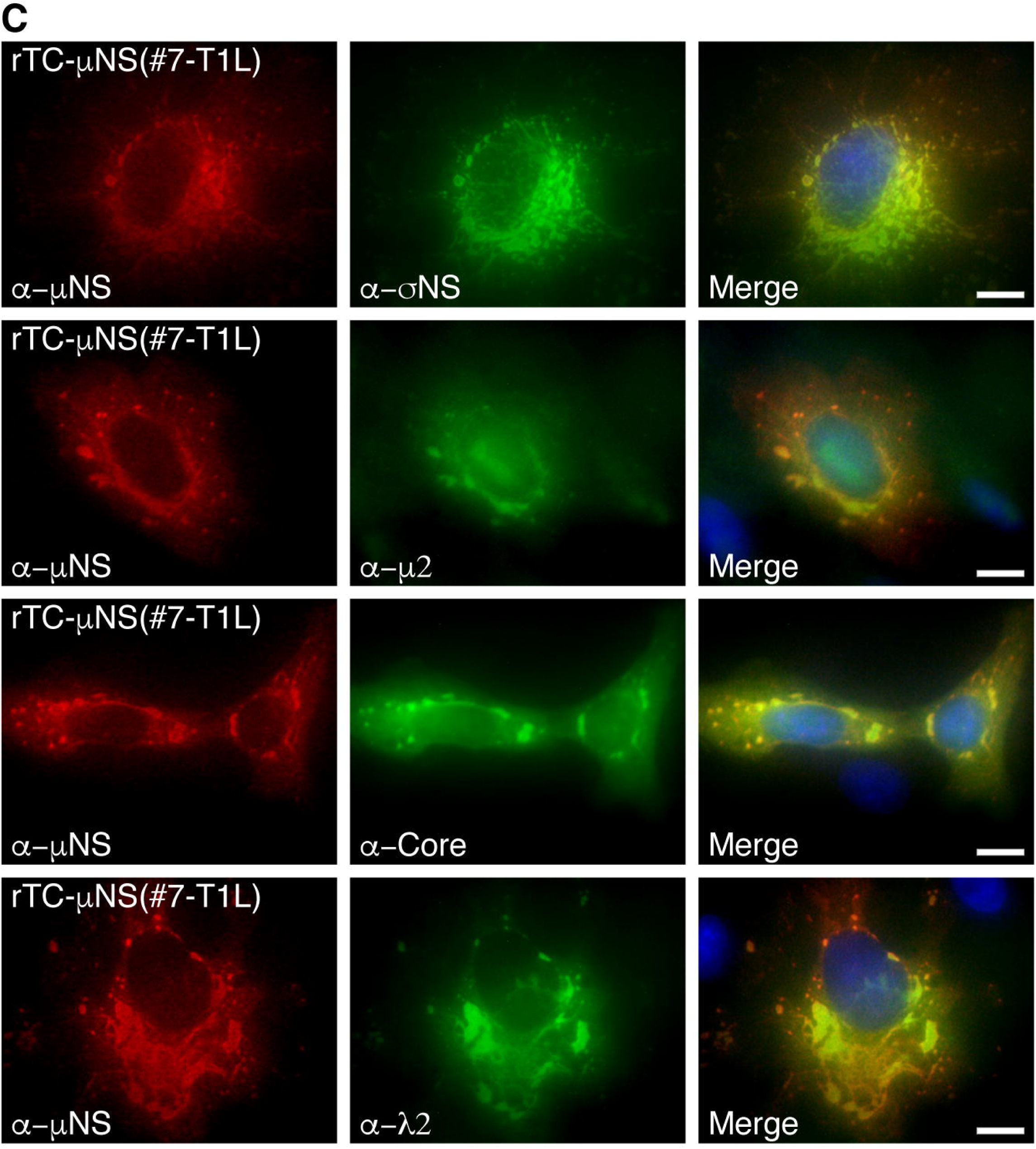
Colocalization of MRV proteins with recombinant virus-expressed μNS. CV-1 cells were infected with (A) T1L, (B) rTC-μNS(C-term-T1L)/P2, or (C) rTC-μNS(#7-T1L)/P2, and at 18 h p.i., immunostained with mouse (second and third rows) or rabbit (first and fourth rows) antibodies against μNS and mouse σNS antibodies (first row), rabbit μ2 antibodies (second row), MRV core rabbit antibodies (third row), λ2 mouse antibodies (fourth row), followed by Alexa 594-conjugated donkey anti-rabbit or anti-mouse IgG (first column) and Alexa 488-conjugated donkey anti-mouse or antirabbit IgG (second column). Merged images with DAPI-stained nuclei are also shown (third column). Bars = 10 μm.

### Recombinant virus growth deficiency occurs after viral entry, transcription and translation

As each recombinant virus was capable of forming VFs and recruiting known μNS associating proteins to VFs similar to wildtype, we further investigated their growth deficiency to identify at which stage of growth the defect occurs. We began by performing plaque assays to determine PFUs of specific stocks of wildtype and rTC-μNS viruses. We then immediately used these PFU calculations and the same viral stocks to perform: 1) immunoblot analysis on μNS and σNS expression throughout infection to determine if the recombinant viruses had measurable differences to wildtype in virus lifecycle steps leading to protein translation, 2) immunblot analysis of these stocks to measure virion proteins present in identical PFUs of wildtype versus rTC-μNS virus, 3) electropherograms on the virus stocks to determine the amount and ratio of genomic RNA present in the same PFU of wildtype versus recombinant virus, and 4) repeat plaque assays with identical PFUs of virus to confirm our original PFU calculation. L929 cells were infected with wildtype T1L, rTC-μNS(C-term-T1L)/P5, or rTC-μNS(#7-T1L)/P5 and samples were collected at 0, 12, 24, and 48 h p.i. and subjected to immunoblot analysis with antibodies against μNS, to examine specific impacts of the TC-tag insertion on μNS expression, and against σNS, to examine expression of a viral protein other than μNS. Antibodies against α-tubulin were also used as a protein loading control (Fig. 5A). We observed that both recombinant viruses expressed substantially higher amounts of μNS and σNS compared to wildtype T1L at each timepoint, suggesting that the TC-tag within μNS does not disrupt μNS protein expression, and that the recombinant viruses were not defective at early stages of infection including virus entry, RNA transcription, and protein translation. Importantly, this data also suggested that substantially more recombinant virions may be required to achieve the same PFU relative to wildtype virus.

**Figure 5.**
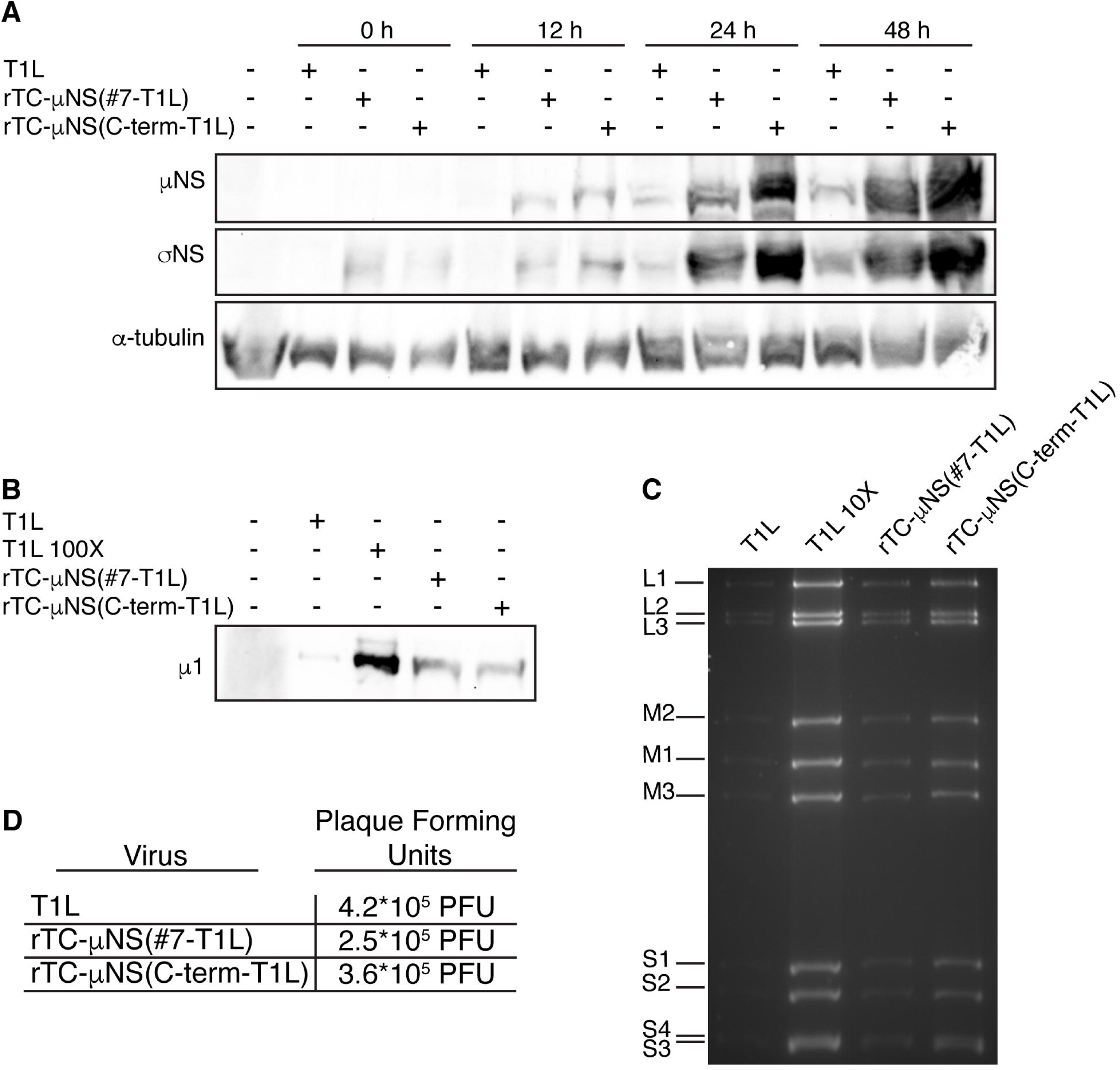
Recombinant virus replication characterization. (A) L929 cells were infected with 5.0×10^5^ PFUs of T1L, rTC-μNS(C-term-T1L)/P5, or rTC-μNS(#7-T1L)/P5 and samples were collected at 0, 12, 24, and 48 h p.i. and were run on SDS-PAGE, transferred to nitrocellulose and immunoblotted with α-μNS (first row), α-σNS (second row), and α-α-tubulin antibodies (third row). (B) 5.0×10^5^ PFUs of T1L, rTC-μNS(C-term-T1L)/P5, and rTC-μNS(#7-T1L)/P5 and 5.0×10^7^ PFUs of T1L (T1L 100X) were run on SDS-PAGE, transferred to nitrocellulose and immunoblotted with T1L α-virion antibody (μ1). (C) RNA extracted from 1×10^8^ PFUs of T1L, rTC-μNS(C-term-T1L)/P5, and rTC-μNS(#7-T1L)/P5 or 1×10^9^ PFUs of T1L (T1L 10X) was run on 10% SDS-PAGE for 12 h to visualize dsRNA genomic segments. (D) 5.0×10^5^ PFUs T1L, rTC-μNS(C-term-T1L)/P5, and rTC-μNS(#7-T1L)/P5 were subject to a plaque assay on L929 cells.

To explore this hypothesis, we examined the virus preparations directly by subjecting identical PFUs of the same wildtype and recombinant virus stocks to immunoblot analysis with antibodies against the virion, which detected substantially more virions/PFU in the recombinant viruses compared to wildtype (Fig. 5B). We additionally examined genomic RNA associated with the same PFUs of wildtype and recombinant virus, and again found that there were substantially higher levels of genome associated with the same PFU of recombinant viruses relative to wildtype (Fig. 5C). Finally, a repeated plaque assay of the same stocks showed that the PFUs between the viruses remained essentially the same (Fig. 5D). Together these findings suggest that the growth defect in the recombinant viruses is downstream of early events such as entry, transcription, and translation, likely at the level of assortment, replication, or assembly. These findings further suggest that the recombinant viruses are capable of producing high levels of infectious virions, and the diminished titers relative to wildtype virus over time (Fig. 3) is likely not a result of a catastrophic replication deficiency, but is instead a result of diminished efficiency in one or more of the later steps in the replication cycle.

### A single recombinant virus retains the TC-tag over ten passages

We next focused our attention on the stability of the TC-tag within the recombinant viruses. Viral RNA was extracted from the second and fifth passage of each recombinant virus using TRIzol LS followed by RT-PCR of the M3 genome segment and sequencing. At passage two, the rTC-μNS(#7-T1L) M3 genome segment showed a mixture of viruses in which the dominant phenotype had a 1649G>A mutation that resulted in a C550Y mutation producing a CCPGCY instead of CCPGCC TC-tag while a smaller portion of the recombinant virus population contained the CCPGCC (Fig. 6A). By passage five the virus had stabilized and exclusively contained the C550Y mutation. This data showed that our earlier FlAsH-EDTA labeling experiments using rTC-μNS(#7-T1L)/P2 contained a mixed population of virus (Fig. 2). As this labeling was diminished relative to rTC-μNS(C-term-T1L), we were curious whether the labeling defect was a result of the C550Y mutation or whether this mutation would lead to a complete loss of labeling. Therefore, we infected CV-1 cells with rTC-μNS(#7-T1L)/P5 which contained the CCPGCY mutation exclusively to determine if μNS would label with FlAsH-EDT2. We found that VF labeling was similarly diminished using this passage, suggesting that the FlAsH-EDT2 reagent was able to maintain binding, albeit to a lesser extent, to the CCPGCY sequence (Fig. 6B).

**Figure 6.**
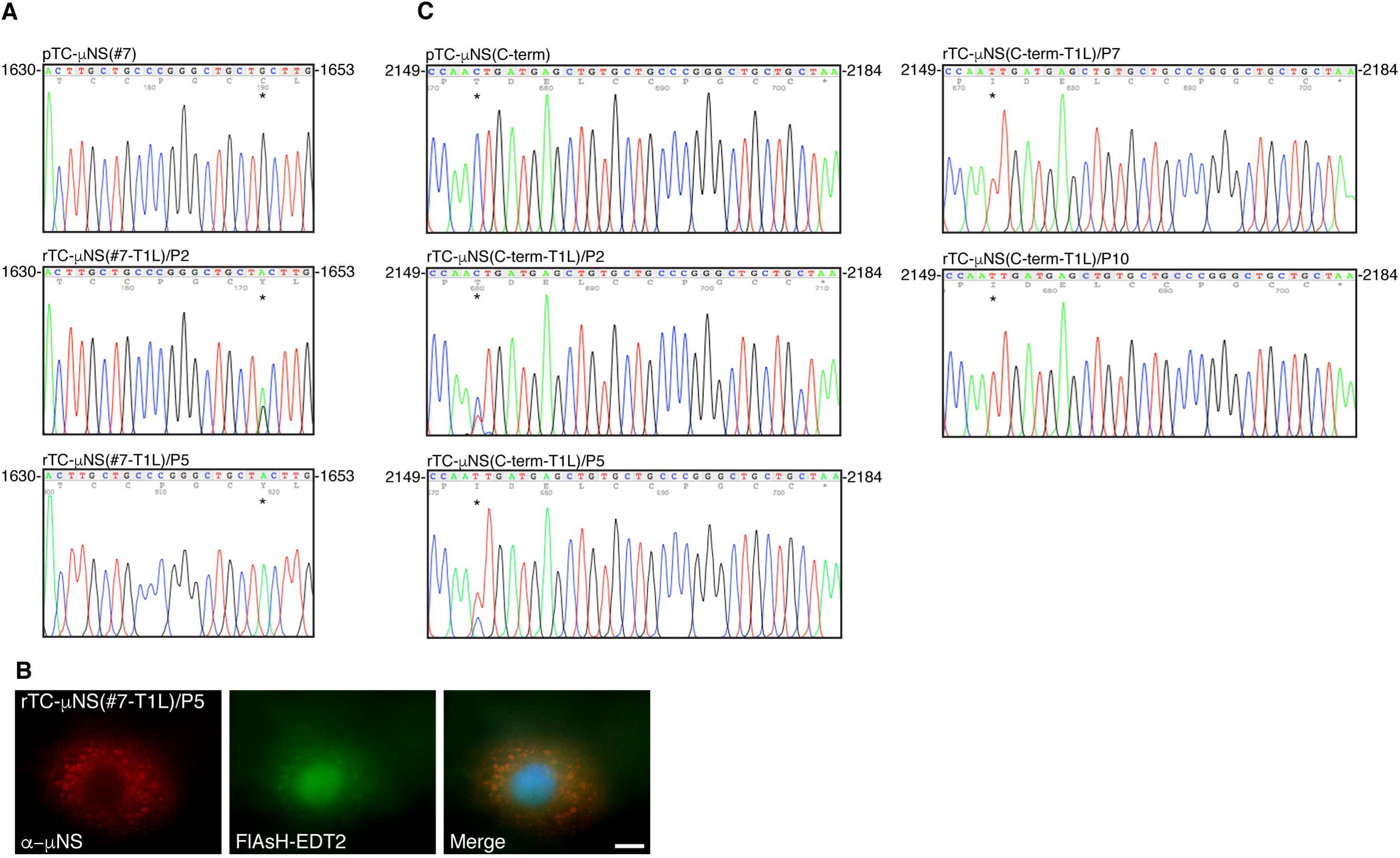
Sequencing of recombinant virus passages. rTC-μNS(C-term-T1L) and rTC-μNS(#7-T1L) viruses were passaged ten times and sequenced to determine the stability of the TC-tag. (A) The pTC-μNS(#7) and rTC-μNS(#7-T1L) passages 2 and 5 were sequenced and the TC-tag and acquired second-site mutation, portrayed with an asterisk (*), are shown. All other regions of M3 were the same as wildtype. (B) CV-1 cells were infected with the rTC-μNS(#7-T1L)/P5 at an MOI of 1, and at 18 h p.i., cells were labeled with FlAsH-EDT2 (middle column) for 45 min, then fixed and immunostained with rabbit α-μNS antibody (left column), followed by Alexa 594-conjugated donkey anti-rabbit IgG. A merged image is also shown with DAPI staining (right column). Images are representative of the observed phenotype. Bar = 10 μm. (C) The pTC-μNS(C-term) and rTC-μNS(C-term-T1L) passages 2, 5, 7, and 10 were sequenced and the TC-tag and acquired second-site mutation, portrayed with an asterisk, are shown. All other regions of M3 were the same as wildtype.

Sequencing the M3 gene of passage two of the rTC-μNS(C-term-T1L) virus also showed a mixed population of viruses in which the CCPGCC TC-tag itself remained stable but the virus had acquired a second-site mutation at 2153C>T, that resulted in a T718I mutation within the protein upstream of the TC-tag (Fig. 6C). This mutation became dominant by passage five. To determine if this was a stable mutation we passaged the rTC-μNS(C-term-T1L) virus five further passages and again sequenced the M3 genome segment at passage seven and ten. We found that the seventh passage virus exclusively contained the T718I mutation and this mutation remained stable at passage ten (Fig. 6C) while both passages retained the original CCPGCC TC-tag. We performed replication assays on wildtype T1L, rTC-μNS(C-term-T1L)/P2, and rTC-μNS(C-term-T1L)/P7 followed by plaque assays to determine if the T718I mutation restored the virus to wildtype T1L replication kinetics or impacted replication in any way, but we observed no significant difference between the two passages (data not shown). Because the rTC-μNS(C-term-T1L) TC-tag remained stable over ten passages and labeled with FlAsH-EDT2 to higher levels than rTC-μNS(#7-T1L), we proceeded forward using the rTC-μNS(C-term-T1L) virus for live cell imaging.

### VF dynamics during MRV infection

To provide a proof of principle that rTC-μNS(C-term-T1L) will be a useful tool to study VF dynamics in infected cells, we performed a short time-lapse live cell experiment where we examined VF movement and interaction at a single time in MRV infection. BHK-T7 cells were infected with rTC-μNS(C-term-T1L)/P2, and at 18 h p.i., FlAsH-EDT2 was added to the cells and incubated for 45 mins. Still images of the infected cells were then captured every 20 seconds for 2 h. Mock infected cells without FlAsH-EDT2, as well as FlAsH-EDT2 labeled cells infected with wildtype T1L at an MOI of 100 were included as controls to demonstrate specificity of the FlAsH-EDT2 reagent (Fig. 7A). At this time point in infection, rTC-μNS(C-term-T1L) was found to form small VFs in the periphery of infected cells that move in short stochastic motions in the cell while larger, more stationary VFs were found at the nuclear periphery (Fig. 7B, Supplementary Mov. 1). Small VFs were highly mobile and could be seen fusing with larger VFs (Fig. 7B: red arrows). In addition, small VFs could be seen moving towards one another or towards larger VFs, touching briefly, and then moving away in a kissing motion (Fig. 7B: yellow arrow) which may be a result of incomplete VF fusion. These data suggest that rTC-μNS(C-term-T1L) will be an extremely useful tool for future in depth studies examining VF dynamics and function during MRV infection.

**Figure 7.**
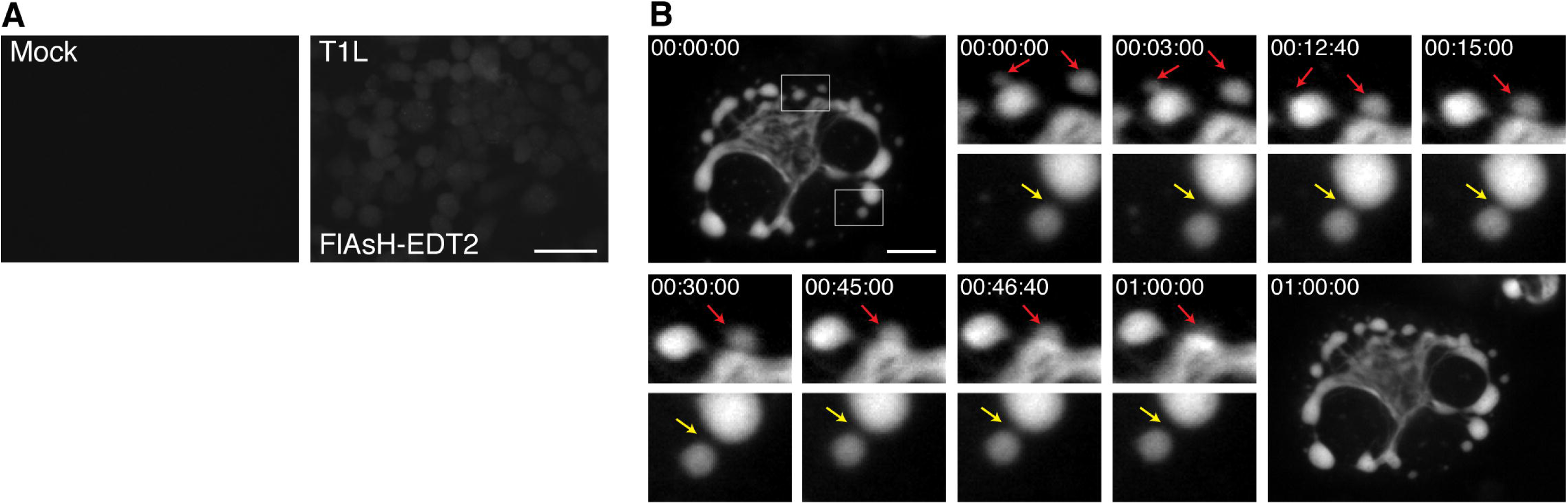
TC-μNS labeled VF dynamics in infected cells. BHK-T7 cells were mock infected or infected with T1L or rTC-μNS(C-term-T1L)/P2, and at 18 h p.i., cells were imaged using live cell microscopy. (A) Mock infected (top left) or T1L infected cells with FlAsH-EDT2 labeling (top right) controls are shown. Bar = 40 μm. (B) rTC-μNS(C-term-T1L)/P2 infected BHK-T7 cell images shown 0-60 mins post FlAsH-EDT2 incubation. Red arrows indicate VF fusion, and yellow arrow indicates kissing motion of VFs (at 4X magnification). Bar = 10 μm.

### VF dynamics in the presence of nocodazole

In addition to facilitating our understanding of VF formation and movement throughout infection, we postulated that rTC-μNS(C-term-T1L) will also be indispensable for understanding the role of VF interactions with cellular proteins throughout infection. One such interaction that has been previously defined is that of μNS and the μNS-interacting protein, μ2, with cellular microtubules (MTs). Destabilization of MTs leads to the formation of small VFs/VFLs concentrated at the periphery of the cell in infected cells or cells transfected with a plasmid expressing μNS, suggesting MTs play a critical role in VF/VFL movement or fusion (9, 33). In addition, the μ2 protein of most MRV strains induces hyperacetylation and stabilization of MTs and also influences the morphology of VFs to a more filamentous, rather than globular VFL phenotype seen when μNS is expressed alone (9, 33). Until recently, most evidence suggested that tight association of VFs with MTs (14) or MT stability (37) in MRV infected cells does not significantly alter viral replication. However, a recent paper has suggested that MT disruption significantly decreases MRV replication by 50%, and further, substantially decreases the crystalline-like array of genome containing virus particles seen in VFs in MRV infected cells with intact MTs, suggesting MTs play a critical role in genome packaging (38). We utilized our rTC-μNS(C-term-T1L) virus to further explore the role of MTs in VF movement and/or fusion within the cell. Vero cells were mock infected or infected with T1L or rTC-μNS(C-term-T1L)/P5 at an MOI of 100 with and without 10 μM nocodazole treatment at 6 h p.i., which destabilizes MTs (Fig. 8A). At 10 h p.i. cells were labeled with FlAsH-EDT2 and at 12 h p.i. FlAsH-EDT2 was removed and still images were taken of VFs every 2 mins for 2 h. In cells not treated with nocodazole we observed stable filamentous VFs that were presumably MT-associated as well as globular VFs which exhibited fluid movement around the cell resulting in some VF fusion events (Fig. 8B, red arrows, Supplementary Mov. 2) similar to those observed in BHK-T7 cells. We also identified a new fusion event in which a small VF moves from the microtubule toward a large VF, fuses, and then pulls the large VF toward the microtubule at which point the large VF fuses with VF material already on the microtubules (Fig. 8B, blue arrows, Supplementary Mov. 2). While this interaction has not previously been identified it might suggest that smaller, more dynamic VFs may be instrumental in assisting and/or facilitating VF fusion. We also recorded disruption of two VFs in close proximity by movement of other VFs directly between them (Supplementary Mov. 2, green arrow).

**Figure 8.**
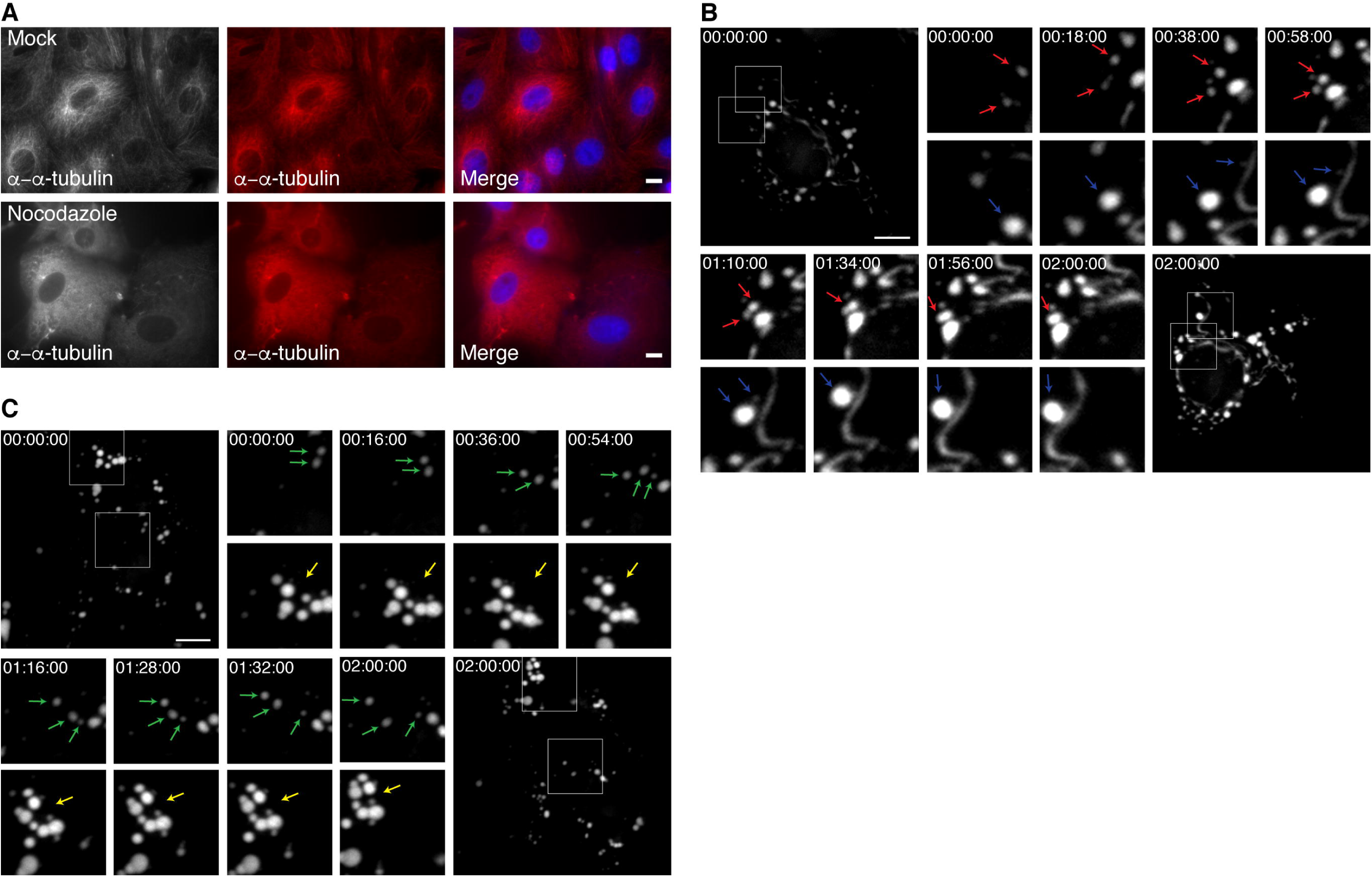
TC-μNS labeled VF dynamics in infected cells with and without nocodazole. Vero cells were infected or mock infected with T1L or rTC-μNS(C-term-T1L)/P5 and at 6 h p.i. cells were treated or untreated with 10 μM nocodazole. (A) At 12 h post treatment, mock infected cells were immunostained with rabbit α-α-tubulin antibody (left and middle columns), followed by Alexa 594-conjugated donkey antirabbit IgG. A merged image is also shown with DAPI staining (right column). (B) At 10 h p.i. T1L and rTC-μNS(C-term-T1L)/P5 infected Vero cells without 10 μM nocodazole were labeled with FlAsH-EDT2 for 2 h at which point rTC-μNS(C-term-T1L)/P5 infected cells were imaged using live cell microscopy. Red and blue arrows indicate VF fusion (at 3X magnification). (C) At 10 h p.i. T1L and rTC-μNS(C-term-T1L)/P5 infected Vero cells with 10 μM nocodazole were labeled with FlAsH-EDT2 for 2 h at which point rTC-μNS(C-term-T1L)/P5 infected cells were imaged using live cell microscopy. The yellow arrow indicates a VF cluster unable to fuse, and green arrows indicate VF movement (at 2X magnification). Bars = 10 μm.

In cells treated with nocodazole, there was no obvious association of VFs with MTs, and all VFs were globular and not filamentous in nature, as expected. Somewhat surprisingly, the VFs appeared to be very dynamic exhibiting short stochastic movements throughout the cell (Fig. 8C, green arrows, Supplementary Mov. 3). While we observed VF movement we did not see accumulation of VFs at the nuclear periphery that we observed in cells without nocodazole treatment. We were also unable to detect any VF fusion events throughout multiple experiments, and instead repeatedly imaged multiple clusters of small VFs that appeared unable to fuse (Fig. 8C, yellow arrow, Supplementary Mov. 3). Together this suggests that the small VF phenotype previously observed following nocodazole treatment in fixed cells (9, 33) is likely a result of an inhibition or inability of VFs to fuse with one another and not a result of total inhibition of VF movement within the cell. In addition to defining a role for MTs in VF fusion and migration, this result further suggests that VFs are able to utilize cellular components apart from microtubules to move within cells.

## Discussion

In this paper we have demonstrated recovery of T1L recombinant MRV with a TC-tag introduced into the non-structural VF matrix protein, μNS. This is the first time a replicating recombinant MRV has been created in which VFs encoded from the genome can be fluorescent labeled during infection. Adding the TC-tag to μNS in a recombinant virus allows for several improvements to existing technologies. Foremost in these improvements is the ability to visualize VFs as they form from a modestly modified μNS protein expressed from virus transcripts. As our rTC-μNS virus undergoes the full MRV replication cycle, findings from future studies using our virus should accurately reflect VF dynamics and interactions throughout MRV infection. In this study, we limited our observation of VFs over short time-lapses, however, there is obvious potential to increase the observation period to a full replication cycle to observe initial formation of VFs and the interactions between VFs throughout the MRV lifecycle. The detection limit of FlAsH-EDT2 labeling of TC-μNS is currently under investigation, however, as we could observe quite small VFs and VFLs in our studies, we expect this approach to be quite useful in delineating early versus late events in VF formation and maturation.

There are several other questions that can be explored using this virus. Because the FlAsH-EDT2 reagent can be removed from cells at any time, pulse chase studies can be performed to examine TC-μNS stability within VFs over time. Moreover, a second reagent, ReAsH-EDT2 (25), which fluoresces in the red spectrum (608 nm) can be used subsequent to FlAsH-EDT2 labeling to determine differences in behavior of existing TC-μNS versus newly translated TC-μNS. In addition, as we demonstrate with MTs and nocodazole, FlAsH-EDT2 labeling can also be used in combination with inhibitory drugs and other fluorescent labeling techniques to examine VF interactions with cellular proteins that may modulate or be modulated by VFs, μNS, or other VF-localized virus proteins. Finally, although it will likely be dependent on location of the TC-tag within individual virus proteins, demonstration of recovery of recombinant viruses containing the small TC-tag in μNS suggests that other MRV proteins may be amenable to TC-tagging, allowing their visualization during infection.

Apart from VF dynamics the study of the TC-μNS mutants and recombinant viruses presented in this study also produced some new insight into requirements for VFL and VF formation and μNS binding with other MRV proteins. One result we found particularly interesting was that we were unable to recover the TC-μNS(#4) mutant which contains the TC-tag within a region of μNS known only to be the site of Hsc70 binding, which is not required for normal VF formation (35), but were able to recover two mutants containing the TC-tag within the C-terminal third which is known to be important for VF formation. Our inability to rescue TC-μNS(#4) as well as the other mutants may be due to protein misfolding, but it is also possible that misfolding of the M3 RNA resulted in improper assortment or packaging thus prohibiting viral recovery. We also found it interesting that the TC-μNS(#7) VFLs had a significantly diminished localization with μ2, λ1, and λ3 compared to wildtype μNS, but visually the rTC-μNS(#7-T1L) VFs localized to associating proteins similar to wildtype. This may suggest that μNS associating protein localization to VFs during infection are facilitated by interactions between multiple proteins and RNA; therefore, a defect in association between μNS and each individual protein is magnified in the absence of the full complement of VF-localized interactions in transfected cells.

During characterization of the recombinant viruses we concluded that both recombinant viruses were likely attenuated downstream of viral entry, transcription, and translation. Further investigation into recombinant VF formation and localization with μNS associating proteins compared to wildtype VFs demonstrated that factory formation also was not considerably altered, suggesting that the recombinant VFs closely mimic natural VFs at least to the stage of infection where prominent VFs are constructed. This points to assortment, assembly, or replication as responsible for the recombinant virus growth attenuation. This could simply be a result of the TC-tag inhibiting a function of the μNS protein downstream of VF formation, or alternatively could be a consequence of the nucleotides encoding the TC-tag disrupting proper M3 mRNA folding, resulting in inhibition of assortment, packaging, or replication of the viral genome. Presumably as a result of this attenuation, the rTC-μNS(#7-T1L) lost the inserted TC-tag sequence within two passages. The rTC-μNS(C-term-T1L) virus maintained the TC-tag over ten passages but instead developed a T718I mutation upstream of the TC-tag, which has not been previously documented in the M3 gene of MRV strains. We initially believed this may be a compensatory mutation, however, when rTC-μNS(C-term-T1L)/P7 was directly tested against rTC-μNS(C-term-T1L)/P2 and wildtype T1L, viral replication remained similar to the earlier passage relative to wildtype (data not shown). Nonetheless, since rTC-μNS(C-term-T1L) maintains the TC-tag, produces VFs that appear identical to wildtype VFs, and has not acquired additional second-site mutations over five subsequent passages, we believe rTC-μNS(C-term-T1L) has already in this study, and will continue to enhance our ability to understand VF formation and function during MRV infection.

While it has already been shown that MRV required MT stability to form large, perinuclear VFs (9, 33), whether MTs are necessary for VFs to move into close proximity in order to fuse or strictly for fusion of VFs was unknown. Our data suggests that even in the presence of the MT destabilizing drug, nocodazole, VFs demonstrate movement within the cell (Fig. 8C, Supplementary Mov. 3). However, without stable MTs, VF fusion is a rare or non-existent event, suggesting that while MT stability is dispensible for VF movement, it is necessary for VF fusion. This also suggests that VFs may utilize other cellular components to move throughout the cell. It has previously been shown that nocodazole treatment results in many small VFs in the periphery of cells, as opposed to the perinuclear space (9, 33). Although our studies suggest that VFs are able to move in the absence of MT stabilization, this movement did not appear to result in a directed migration of the non-fused VF clusters toward the nucleus. Instead VF movement was limited to small changes in location independent of direction, indicating that while VFs can move and appear to associate with each other in cells without MTs, directed overall migration of VFs to the perinuclear space appears to require MT stabilization. Finally, our data also shows that once globular VFs associate with MTs to form filamentous VFs, they appear to remain associated with them and become relatively static and stable compared to globular VFs. Taken together with published data, our findings suggest that the path to mature VF formation includes steps where, 1) μNS first self-associates and/or associates with other viral and cellular proteins to form small globular VFs, 2) these small globular VFs move independent of MTs in the cell and encounter each other, 3) following this encounter, smaller VFs fuse into larger VFs in a MT-dependent manner or fuse to filamentous VFs already associated with MTs, and 4) the MT-associated VFs then either migrate towards, or accumulate in the perinuclear space within the cell where they remain relatively stable over time.

## Acknowledgments

We thank Dr. Terry Dermody for the MRV reverse genetics plasmids and Dr. Pooja Gupta-Saraf for technical assistance. This work was supported by the Bailey Research Career Development Award to C.L.M. as well as the F. Wendell Miller Fellowship to L.D.B. Other assistance to C.L.M. was provided by the Office of the Dean, Iowa State University College of Veterinary Medicine.

